# Metformin impairs trophoblast metabolism and differentiation in dose dependent manner

**DOI:** 10.1101/2023.02.14.528531

**Authors:** Sereen K. Nashif, Renee M. Mahr, Snehalata Jena, Seokwon Jo, Alisa B. Nelson, Danielle Sadowski, Peter A. Crawford, Patrycja Puchalska, Emilyn U. Alejandro, Micah D. Gearhart, Sarah A. Wernimont

## Abstract

Metformin is a widely prescribed medication whose mechanism of action is not completely defined and whose role in gestational diabetes management remains controversial. In addition to increasing risks of fetal growth abnormalities and preeclampsia, gestational diabetes is associated with abnormalities in placental development including impairments in trophoblast differentiation. Given that metformin impacts cellular differentiation events in other systems, we assessed metformin’s impact on trophoblast metabolism and differentiation. Using established cell culture models of trophoblast differentiation, oxygen consumption rates and relative metabolite abundance were determined following 200 μM (therapeutic range) and 2000 μM (supra-therapeutic range) metformin treatment using Seahorse and mass-spectrometry approaches. While no differences in oxygen consumption rates or relative metabolite abundance were detected between vehicle and 200 μM metformin treated cells, 2000 μM metformin impaired oxidative metabolism and increased abundance of lactate and TCA cycle intermediates, α-ketoglutarate, succinate, and malate. Examining differentiation, treatment with 2000 μM, but not 200 μM metformin, impaired HCG production and expression of multiple trophoblast differentiation markers. Overall, this work suggests that supra-therapeutic concentrations of metformin impairs trophoblast metabolism and differentiation whereas metformin concentrations in the therapeutic range do not strongly impact these processes.

## Introduction

Metformin (1,1-dimethylbiguanide) is a first-line treatment for patients with type 2 diabetes given its ability to improve glucose levels, support weight loss, and reduce cardiovascular morbidity (Maruthur et al., 2016). Until recently, metformin was also considered a first-line medication for gestational diabetes mellitus (GDM), improving glucose levels, maternal weight gain, and preeclampsia (Liang et al., 2017;2018;D’Ambrosio et al., 20l9;ElSayed et al., 2023). However, metformin is no longer considered a first line medication for GDM since long term follow up has demonstrated increased weight, body mass index, and adiposity at mid-childhood following in utero metformin exposure (Tarry-Adkins et al., 2019). Despite these concerns, there continues to be interest in use of metformin in pregnancy for potential preeclampsia prevention, reduced maternal weight gain in obese pregnancies, and lower miscarriage rate among pregnancies complicated by polycystic ovarian syndrome (Syngelaki et al., 2016;Løvvik et al., 2019;Adams et al., 2022). However, the mechanisms underlying these potential beneficial outcomes remain unknown.

During pregnancy, the placenta grows to support the health of the developing fetus (Turco and Moffett, 2019). A key cell facilitating nutrient transport to the developing fetus is the syncytiotrophoblast, a unique multi-nucleated cell that supports nutrient transport and gas exchange between maternal and fetal circulation (Lager and Powell, 2012). During the course of pregnancy, cytotrophoblasts divide and fuse to support the development and health of the syncytiotrophoblast (Costa, 2016;Renaud and Jeyarajah, 2022). Trophoblast differentiation is critical for placental development and fetal health. Delayed villous maturation, the most common histopathologic finding in GDM, is associated with poor birth outcomes (Higgins et al., 2011;Treacy et al., 2013;Huynh et al., 2015;Khong et al., 2016;Jaiman et al., 2020). A key finding in delayed villous maturation is an increase in the ratio of cytotrophoblasts to syncytiotrophoblasts, suggestive of potential defects in trophoblast differentiation (Khong et al., 2016). In one limited study, metformin treatment was found to normalize placental histology in pregnancies complicated by GDM (Arshad et al., 2016). However, it is unclear if this is due to improved blood glucose levels or direct impact of metformin on placental development.

Despite its wide clinical use, metformin’s mechanisms of action are not fully understood, and it may have unique impacts within distinct tissues (Rena et al., 2017;Aye et al., 2022). Metformin inhibits complex I of the electron transport chain in a dose dependent manner leading to changes in nucleotide energy charges: this decrease in ATP to ADP and ATP to AMP ratios leads to activation of AMPK (LaMoia and Shulman, 2021). However, AMPK independent mechanisms are also postulated to contribute to improved glycemic profiles (Rena et al., 2017). Impacts of metformin on complex I are demonstrated with supra-therapeutic concentrations, in the 1 to 5 mM range (LaMoia and Shulman, 2021). In contrast, following oral administration, metformin tissue levels are estimated to achieve levels of 50-100 μM (LaMoia and Shulman, 2021). Recent work suggests that maternal and placental metformin concentrations from GDM patients at the time of delivery may be even lower (Tarry-Adkins et al., 2022a). While the timing of metformin administration relative to blood collection is not clear in this study, maternal serum levels are reported in the 0-5 μM range and are linearly correlated to placental metformin levels. This is significantly lower than the 1-5 mM range thought to directly inhibit complex I, suggesting other mechanisms may underlie metformin’s impact in pregnancy (Aye et al., 2022).

Mitochondria regulate cellular differentiation in multiple systems and metformin has been found to promote differentiation in some systems and inhibit it in others (Folmes and Terzic, 2016;Lisowski et al., 2018;Jiang and Liu, 2020;Chakrabarty and Chandel, 2021). Specifically, metformin promotes myogenic, neuronal and osteogenic differentiation through impacts on AMPK (Wang et al., 2012;Dadwal et al., 2015;Senesi et al., 2016). Conversely, metformin inhibits differentiation of cancer stem cells in gastric cancer, osteosarcoma, and breast cancers (Cheung et al., 2019;Tan et al., 2019;Zhao et al., 2019). The impact of metformin on trophoblast differentiation has not been reported.

Given the ongoing interest in metformin for the clinical management of obstetric complications and its known impact on cellular differentiation in other systems, we hypothesized that metformin regulates trophoblast differentiation and cellular metabolism. In this work, we employ cell culture models to test the impact of metformin at near therapeutic and supra-therapeutic concentrations on trophoblast metabolism and differentiation. While supra-therapeutic doses of metformin impair cellular metabolism and trophoblast differentiation, no significant impact was seen at therapeutic concentrations.

## Materials and methods

### Ethics

This study does not meet criteria for human subjects research, confirmed by the University of Minnesota Institutional Review Board (HRD # STUDY00014203).

### Cell Culture and Reagents

BeWo cells were purchased from ATCC and maintained in culture with equal parts F12K (ATCC) (supplemented with 10% Fetal Bovine Serum (R&D Systems), Penicillin-Streptomycin (Gibco)) and DMEM Complete (4.5g/L glucose, Gibco) (supplemented with Glutamine (Gibco), Sodium Pyruvate (Gibco), 10% Fetal Bovine Serum (Gibco), and Penicillin-Streptomycin (Gibco)). To induce syncytialization, BeWo cells were cultured on tissue culture treated plates for 24 hours. Forskolin (Millipore Sigma) (40 μM) was added for 48 hours with DMSO (0.4%) (Sigma-Aldrich) as the vehicle control. Metformin (Cayman) at concentrations of 200 μM or 2000 μM was added for 72 hours. PBS (Gibco) was used as a vehicle control for metformin. Media and treatments were replaced every 24 hours during syncytialization.

Trophoblast Stem Cells (TSC) (CT29, Acquired from RIKEN Cell Bank) were maintained in a self-renewing state on iMatrix511 (Amsbio) coated plates in TS Complete media. TS Complete media is comprised of DMEM/F12 media (Gibco) containing 100 μg/ml primocin, 0.15% BSA, 1% ITS-X, 1% Knockout Serum Replacement and 0.2 mM ascorbic acid and supplemented with 2.5 μM Y27632, 25 ng/ml EGF, 0.8 mM Valproic Acid, 5 μM A83-01, and 2 μM CHIR99021 (Okae et al., 2018). Cells were cultured at 37°C, 5% CO_2_. On day 0, cells were plated on a 6 well plate coated with iMatrix511 in TS Complete media and metformin (Cayman) at concentrations of 200 μM or 2000 μM. PBS (Gibco) was used as a vehicle control. On day 1, to induce syncytialization, TS Complete media was replaced with ST Differentiation media containing DMEM/F12 with 100 μg/ml Primocin, 0.1% BSA, 1% ITS-X, 2.5 μM Y27632, 4% KSR, and 2 μM forskolin. Metformin at the respective concentrations was also added during the media change. Self-renewing cells were maintained in TS Complete media with metformin or PBS. On day 4 after plating, RNA was isolated for qPCR.

### Protein Expression

Cells were washed with ice cold PBS and lysed with RIPA Buffer (Sigma) supplemented with Protease Inhibitors (Thermo Fisher Scientific), phosphatase inhibitor cocktail (Pierce) and 1mM DTT (Sigma) in a 6 well plate. Lysate was then centrifuged for 10 minutes at 4 C at 21,000X*g*. Pierce Bicinchoninic Acid Protein Assay Kit (Thermo Scientific) was used to determine protein concentration. 30 μg of protein per well was used to detect protein expression levels. Lysates were run on 4-12% Bis-Tris gel (Invitrogen), transferred to PVDF, and blocked with 5% milk (Research Products International). Blots were stained overnight at 4 C with total AMPK (Cell Signaling Technology) and pAMPK Thr 172 (Cell Signaling Technology) antibodies in 2% bovine serum albumin (Gemini Bio). Blots were washed with Tris-Buffered Saline and stained with peroxidase conjugated secondary antibodies (Jackson ImmunoResearch). Blots were developed using SuperSignal West Pico PLUS Chemiluminescent Substrate (Thermo). Total protein loading was assessed by BlotFastStain (G-Biosciences). Blots were imaged using Bio-Rad ChemiDoc MP imaging system and band density was quantified using ImageJ.

### Seahorse Assay

30,000 BeWo cells were plated into wells of Cell-Tak-coated XFe96 plates containing 100 μL/well of equal parts F12K (ATCC) and DMEM (Gibco). Cells were plated in the presence of PBS, 200 μM metformin, or 2000 μM metformin, and treated with DMSO or 40 μM forskolin 24 hours after plating. 72 hours after plating, Seahorse XF cell Mito stress test was performed to measure ECAR and OCR using the Seahorse Extracellular Flux (XFe96) Analyzer (Seahorse Bioscience Inc. North Billerica, MA). On the day of the assay, the media was switched to 180 μL of Seahorse XF assay media (DMEM) freshly supplemented with 16 mM glucose, 10 mM sodium pyruvate, and 2 mM glutamine (Agilent, Santa Clara, CA). The plate was incubated in a 37°C non-CO_2_ incubator for 1h prior to measurement. The plate was then transferred to the Seahorse XFe96 Analyzer (Seahorse Bioscience Inc. North Billerica, MA) for analysis. Once in the instrument, cells underwent measurement of basal oxygen consumption and extracellular acidification, followed by successive treatments with oligomycin A (2μM), FCCP (carbonyl cyanide-p-trifluoromethoxyphenylhydrazone) (1μM), and rotenone and antimycin A (0.5 μM). OCR and ECAR measurements were normalized per well to DNA, measured using the Quant-iT PicoGreen dsDNA Assay kit. For analysis, post-rotenone/antimycin A OCR readings were subtracted from rest of OCR values to set baseline mitochondrial respiration. Key bioenergetic parameters (basal respiration, ATP linked respiration, maximal respiration, spare capacity, and proton leak) from the mito stress test were calculated according to manufacturer’s protocol **(Figure 2B).**

### Metabolomics

250,000 BeWo cells were plated on 6 well plates, treated with vehicle or metformin and 24 hours after plating, treated with DMSO or 40 μM forskolin as above. A mirror plate was made to facilitate normalization of metabolite abundance to total protein. 72 hours after plating, samples were collected by washing twice with ice cold PBS, once in ice cold water, and snap freezing in liquid nitrogen prior to transferring cells in methanol to a fresh tube. Solvent was evaporated and samples were stored at −80 C until ready for metabolite extraction and analysis.

#### Relative Abundance of glycolytic and TCA cycle intermediates

To determine relative abundance of selected TCA cycle and glycolytic metabolites, metabolites were extracted as previously described (Ivanisevic et al., 2013;Puchalska et al., 2018;Puchalska et al., 2019), with modifications. Briefly, 1000 μL of 2:2:1 Acetonitrile (AcN):Water(H_2_0):MeOH (v:v:v) was added to each sample, which underwent three rounds of vortexing, sonication and snap freezing in liquid nitrogen. The samples were incubated at −20°C for 1-4 hours, spun to remove proteins, transferred to fresh tubes, evaporated, and reconstituted in 40 μL of 1:1 AcN:H_2_O. All analyses were performed on Thermo Vanquish liquid chromatograph with Thermo Q-Exactive Plus mass spectrometer equipped with heated ESI (HESI) source.

Samples were analyzed using negative mode on Atlantis Premier BEH Z-HILIC Column (2.1 mm × 100 mm, 1,7 μm). Separation was complete using gradients of Mobile phase A (15 mM ammonium bicarbonate in water, pH 9.0) and Mobile Phase B (15 mM ammonium bicarbonate pH, 9.0 with 90% AcN). Binary gradients of 10% Mobile Phase A for 5 min, 35% Mobile Phase A for 2 min and 10% Mobile Phase A for 3 minutes. Separations were performed at a flow rate 0.5 to 1 mL/min and column temperature at 30°C with an injection volume of 2 μL. The mass spectrometer was operated in negative mode using full scan (FS) mode *(m/z* 68–1,020) with optimized HESI source conditions: auxiliary gas 10, sweep gas 1, sheath gas flow at 30 (arbitrary unit), spray voltage −4 kV, capillary temperature 350°C, S-lens RF 50, and auxiliary gas temperature 350°C. The automatic gain control (AGC) target was set at 3e6 ions and resolution was 70,000.

Xcalibur’s QuanBrowser from Thermo was used for peak identification and integration. Metabolite profiling data was analyzed using verified peaks and retention times. Metabolite peak identity was confirmed based on retention time, *m/z,* and compared to authentic standards as described previously (Puchalska et al., 2018;Puchalska et al., 2019). Integrated signal for each metabolite was normalized to total protein determined using Bicinchoninic Acid Assay (BCA) on mirror plates for each condition.

### HCG production

After treatment, media was collected and samples were tested for alpha HCG using ELISA assay (DRG International). DMSO treated samples were diluted (1:2) with PBS and Forskolin treated samples were diluted (1:100). HCG expression was detected using the Gen5 program via the Synergy HTX Multimode Reader (BioTek).

### Analysis of mRNA expression

Cells were washed with ice cold PBS. Total RNA was extracted from cells using the Quick RNA Mini-Prep extraction Kit (Zymo Research). Equal amounts of RNA were used to synthesize cDNA (BioRad). For quantitative real time PCR, reactions were carried out using SsoAdvanced Universal SYBR Green Supermix (BioRad) on a CFX384 Real-Time System (Bio-Rad). Transcripts were quantified using the 2^-ΔΔCT method.^ and normalized to cyclophilin. See key resource table for specific primers.

### Imaging

50,000 BeWo cells were plated on poly-L-lysine pre-treated glass coverslips along with media containing PBS vehicle or 200 μM or 2000 μM metformin. 24 and 48 hours after plating, DMSO or 40 μM forskolin was applied. 72 hours after plating coverslips were fixed with 4% paraformaldehyde (Fisher Scientific) for 10 minutes and then permeablized using 0.1% TritonX for 10 minutes. After washing, coverslips were blocked for 1 hour with 5% goat serum in PBS. Coverslips were stained overnight at 4 C with HCG (Abcam) and eCadherin (BD Biosciences) antibodies. The following day, coverslips were washed three times, then incubated at room temperature with mouse (Jackson ImmunoResearch) and rabbit (Jackson ImmunoResearch) secondary antibodies. Coverslips were again washed and mounted on glass slides using mounting media with Prolong Glass Antifade Mountant with NucBlue nuclear Stain (Invitrogen). Images were acquired on an Olympus FluoView BX2 Upright Confocal microscope at the University of Minnesota Imaging Core. Fusion index was quantified as the number of nuclei in syncytia divided by the total number of nuclei using Fiji.

### Quantification and Statistical Analysis

Statistical outliers were excluded following identification by the Grubbs test. ANOVA was used to compare the means of more than two groups. *p* values less than 0.05 were considered statistically significant. Data were analyzed using GraphPad Prism.

## Results

### Metformin treatment increases pAMPK signaling in BeWo cells

The BeWo cell line is an established model system where forskolin, an adenylase cyclase activator upstream of protein kinase A, induces both biochemical and morphologic syncytialization (Orendi et al., 2011;Rothbauer et al., 2017). These differentiation parameters are detected through production of human chorionic gonadotrophin (HCG) and coordinated changes in gene transcription along with fusion of cell membranes resulting in morphologic syncytia (Costa, 2016;Renaud and Jeyarajah, 2022).

Using the BeWo model system, we first validated metformin activity within undifferentiated cells by assessing AMP-activated protein kinase (AMPK) activity. Metformin increases AMPK activity through phosphorylation of Thr172 in the activation loop of the kinase domain (Musi and Goodyear, 2002). Following treatment of BeWo cells with vehicle (PBS), 200 μM metformin, or 2000 μM metformin for 72 hours, phosphorylation of AMP-activated protein kinase (pAMPK at Thr172) relative to total AMPK was detected by western blotting **(Figure 1A,B).** Upon treatment with 200 μM metformin, a trending but statistically insignificant increase in pAMPK at Thr172 was observed. However, 2000 μM metformin, increased pAMPK at Thr172 compared to total AMPK. This overall suggests that metformin enters BeWo cells and increases AMPK activity at the 2000 μM concentration.

**Figure 1:**
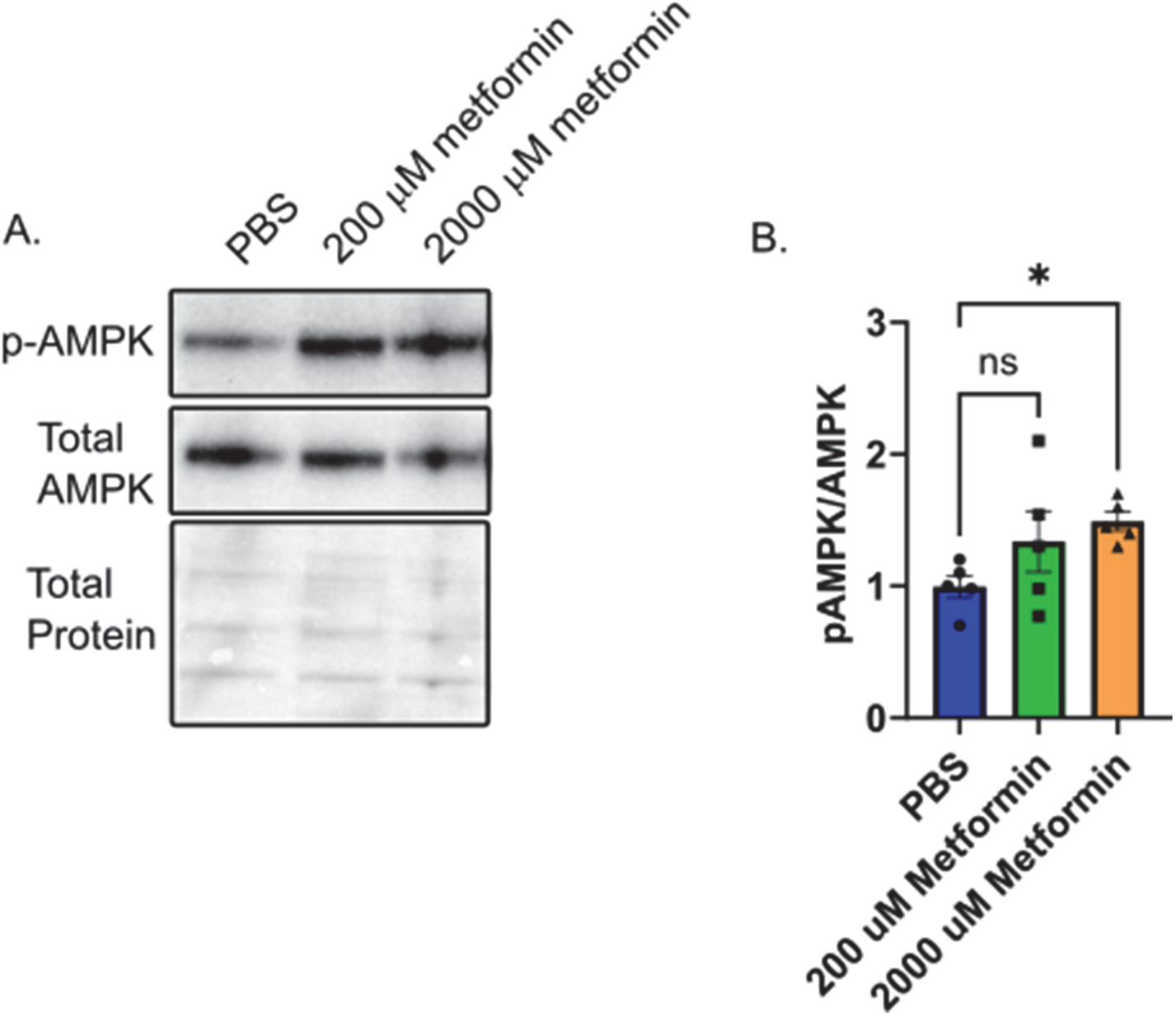
Metformin treatment increases pAMPK signaling in BeWo cells. A) Representative western blot of HCG, pAMPK, and Total AMPK in BeWo cells treated with two different concentrations of Metformin (200 μM or 2000 μM) compared to vehicle (PBS). B) Quantification of Western blots (n=5) demonstrating relative expression of pAMPK to total AMPK in BeWo cells treated with two different concentrations of Metformin (200 μM or 2000 μM) compared to vehicle. Data are representative of mean +/- SEM. *, *p*<0.05.

### Metformin’s Impact on Oxidative and Glycolytic Metabolism

Prior work demonstrates that cytotrophoblasts and syncytiotrophoblasts differ in their glycolytic and oxidative metabolism (Kolahi et al., 2017;Bucher et al., 2021). Given this, we tested the impact of metformin on undifferentiated and differentiated trophoblast metabolism. BeWo cells were plated in the presence of vehicle (PBS), 200 μM or 2000 μM metformin. 48 hours after inducing differentiation with forskolin, a Seahorse Mitochondrial Stress Assay was performed to measure how metformin impacts oxygen consumption rate (OCR) following sequential addition of inhibitors as depicted in **Figure 2 A,B.** Forskolin treatment decreases basal respiration, ATP-linked respiration, maximum respiration, and spare capacity compared to DMSO treated control cells **(Figure 2 C-F).** However, no differentiation dependent change in proton leak or non-mitochondrial respiration was detected. Differentiation in the presence of 200 μM metformin similarly decreased forskolin-dependent basal respiration, ATP-linked respiration, maximum respiration, and spare capacity with no differences detected between vehicle and 200 μM metformin OCR. In contrast, differentiation in the presence of 2000 μM metformin profoundly decreased basal, ATP-linked, maximal, and non-mitochondrial respiration compared to vehicle treatment. Overall, this demonstrates that 2000 μM metformin, not 200 μM metformin, has broad impact on oxidative metabolism **(Figure 2A, C-H).**

**Figure 2:**
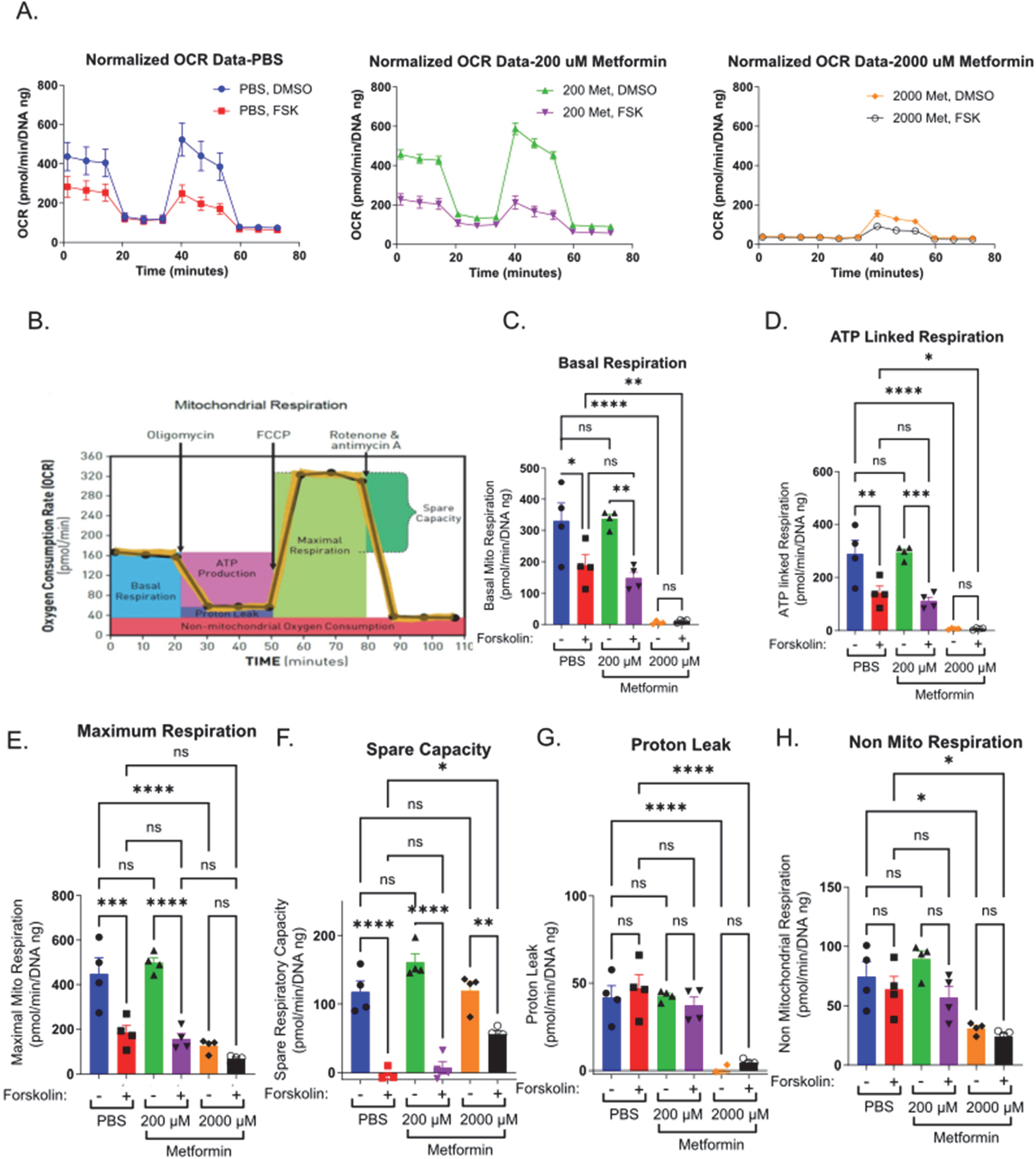
Metformin impairs oxygen consumption rate at high concentrations. A) Normalized Oxygen Consumption Rate (pmol/min/DNA ng) during mitochondrial flux assay for BeWo cells treated with vehicle, 200 μM metformin, or 2000 μM metformin in the presence of DMSO (0.4%, vehicle) or 40 μM forsksolin (FSK) for 48 hours. B) Schematic depicting the impact of sequential inhibitor additions to oxygen consumption rate during Seahorse Mitochondria Stress Test and bioenergetic parameters calculated from the assay. C-H) Bar graphs representing C) Basal Mitochondrial Respiration, D) ATP Linked Respiration, E) Maximal Respiration, F) Spare Capacity, G) Proton Leak and H) non-mitochondrial respiration (all in (pmol/min/DNA ng)) for vehicle, 200 μM and 2000 μM metformin treated BeWo cells. n = 4 biologic replicates. Data are representative of mean +/- SEM. *, *p*<0.05; **, *p*<0.01; ***, *p*<0.001; and ****, *p*<0.0001.

Extracellular acidification rate (ECAR) serves as a surrogate measure of glycolysis. Given the profound impacts of 2000 μM metformin on oxidative metabolism, we determined the impact of metformin on basal ECAR. Forskolin treatment slightly increased basal ECAR rates in PBS and 200 μM metformin treated cells, though no statistically significant differences were detected between groups **(Supplemental Figure 1 A, B, D)**. Following treatment with 2000 μM metformin, ECAR increased significantly in DMSO treated cells compared to vehicle **(Supplemental Figure 1 C, D).** This is consistent with higher metformin concentrations impairing oxidative phosphorylation and increasing glycolytic rate.

### Metformin’s impact on relative abundance of glycolytic and TCA cycle intermediates

Given broad differences observed in OCR with differentiation, we next determined how therapeutic and supra-therapeutic concentrations of metformin impact relative abundance of cellular metabolites during differentiation. Using a high-resolution liquid chromatography, mass-spectrometry (LC-MS) approach, we determined relative metabolite abundance following treatment of BeWo cells with DMSO or forskolin in the presence of PBS, 200 μM or 2000 μM metformin. Following differentiation, ATP increased in PBS and 200 μM metformin treated cells **(Figure 3A).** Surprisingly, 2000 μM metformin maintained ATP levels despite the profound impairments found in oxidative phosphorylation **(Figure 2),** though no differentiation dependent increase was found. (**Figure 3A**). This suggests that upon treatment with 2000 μM metformin, increased glycolytic activity may preserve overall ATP abundance.

**Figure 3:**
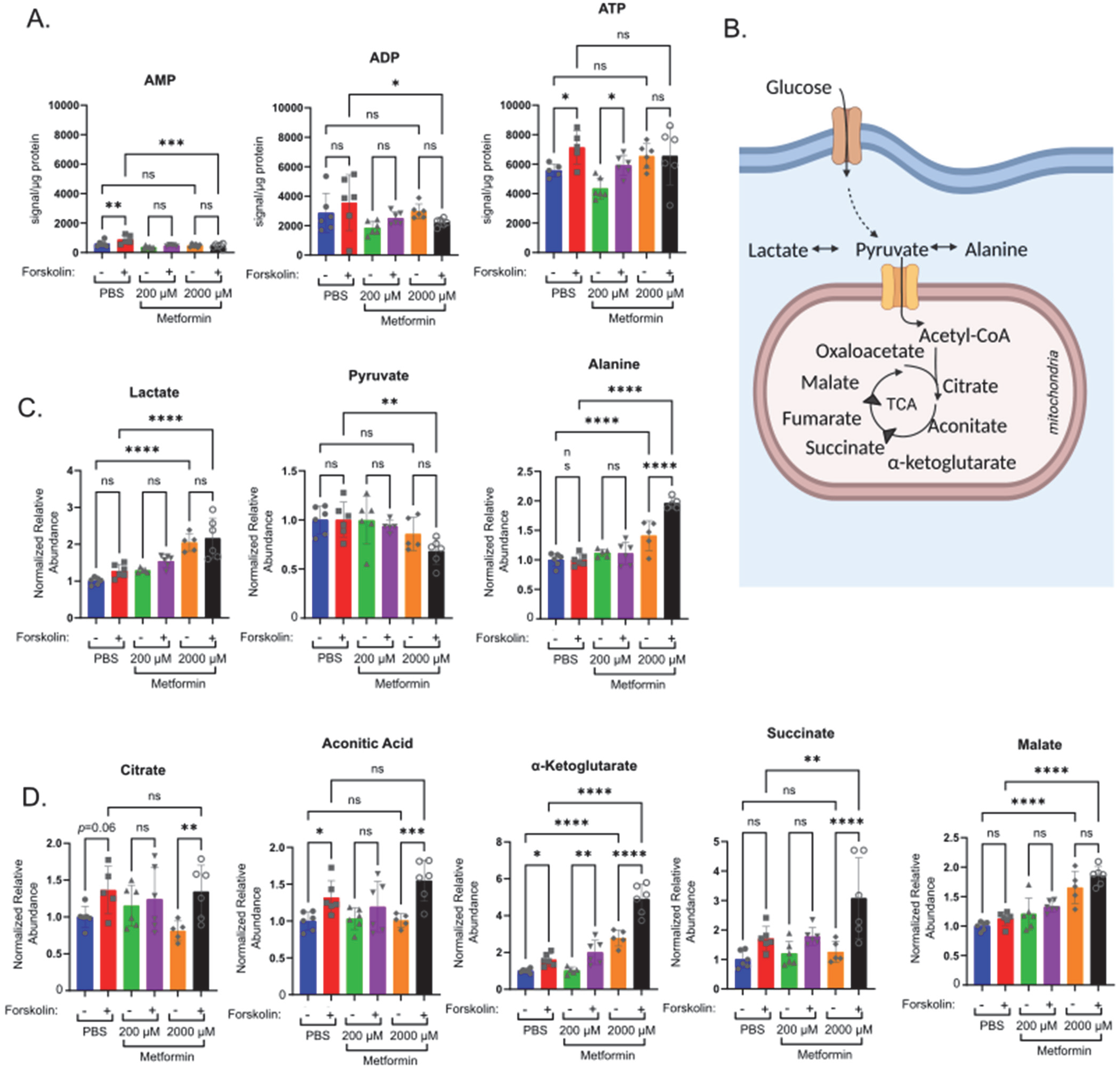
Metformin impacts relative abundance of high energy and TCA cycle metabolites. Forty-eight hours following treatment of BeWo cells with DMSO or Forskolin in the presence of vehicle, Metformin 200 μM, or Metformin 2000 μM, high resolution mass-spectrometry was used to determine relative abundance of metabolites: A) Normalized abundance of AMP, ADP and ATP. Data presented are total metabolite signal normalized to μg protein for each condition. n=6 biologic replicates. B) Schematic demonstrating glucose metabolism to pyruvate, lactate, alanine and TCA cycle metabolites. C) Normalized relative abundance of lactate, pyruvate, and alanine. Normalized relative abundance is total metabolite signal divided by μg protein for each condition and normalized to DMSO treated vehicle control. n=6 biologic replicates. D) Normalized relative abundance of citrate, aconitate, α-ketoglutarate, succinate and malate. Data presented are total metabolite signal divided by μg protein for each condition and normalized to DMSO treated vehicle control. Data are representative of mean +/- SEM. n=6 biologic replicates. *, *p*<0.05; **, *p*<0.01; ***, *p*<0.001; and ****, *p*<0.0001.

We next examined the impact of metformin on lactate, pyruvate, alanine and select TCA cycle intermediates **(Figure 3 B-D).** After glucose enters a cell, it undergoes glycolysis resulting in pyruvate which may be metabolized to lactate, alanine or enter the TCA cycle. Treatment with 2000 μM metformin increased lactate and alanine abundance compared to PBS and 200 μM metformin treatments **(Figure 3C)**. This is consistent with the observed increase in glycolytic metabolism following 2000 μM metformin treatment.

Differentiation in PBS treated cells increased citrate, aconitic acid, α-ketoglutarate, and succinate **(Figure 3D).** This trend persisted in 200 μM treated cells though a differentiation dependent increase was only statistically significant for α-ketoglutarate **(Figure 3D).** In comparison, differentiation following treatment with 2000 μM metformin significantly increased citrate, aconitic acid, α-ketoglutarate, and succinate (**Figure 3D**). This differentiation dependent increase in a-ketoglutarate, succinate and malate with 2000 μM metformin treatment exceeds that observed following vehicle treatment. This suggests that inhibition of oxidative phosphorylation by supra-therapeutic concentrations of metformin may result in accumulation of TCA intermediate metabolites due to impairments in oxidative metabolism.

### Impact of metformin on trophoblast differentiation

Mitochondria regulate cellular differentiation events across multiple systems (Folmes and Terzic, 2016;Lisowski et al., 2018;Chakrabarty and Chandel, 2021). To test the impact of metformin on trophoblast differentiation, we plated BeWo in the presence of vehicle, 200 μM or 2000 μM metformin and treated these cells 24 hours later with vehicle (DMSO) or forskolin to induce syncytialization. We first assessed the impact of metformin on differentiation by assessing HCG production in media samples. HCG is a peptide hormone made by syncytiotrophoblasts (Fournier et al., 2015). We found significantly lower levels of HCG in media samples obtained from cells treated with 2000 μM metformin compared to those in the vehicle control and 200 μM metformin groups **(Figure 4A)**, suggestive of differentiation defects with supra-therapeutic metformin treatment.

**Figure 4:**
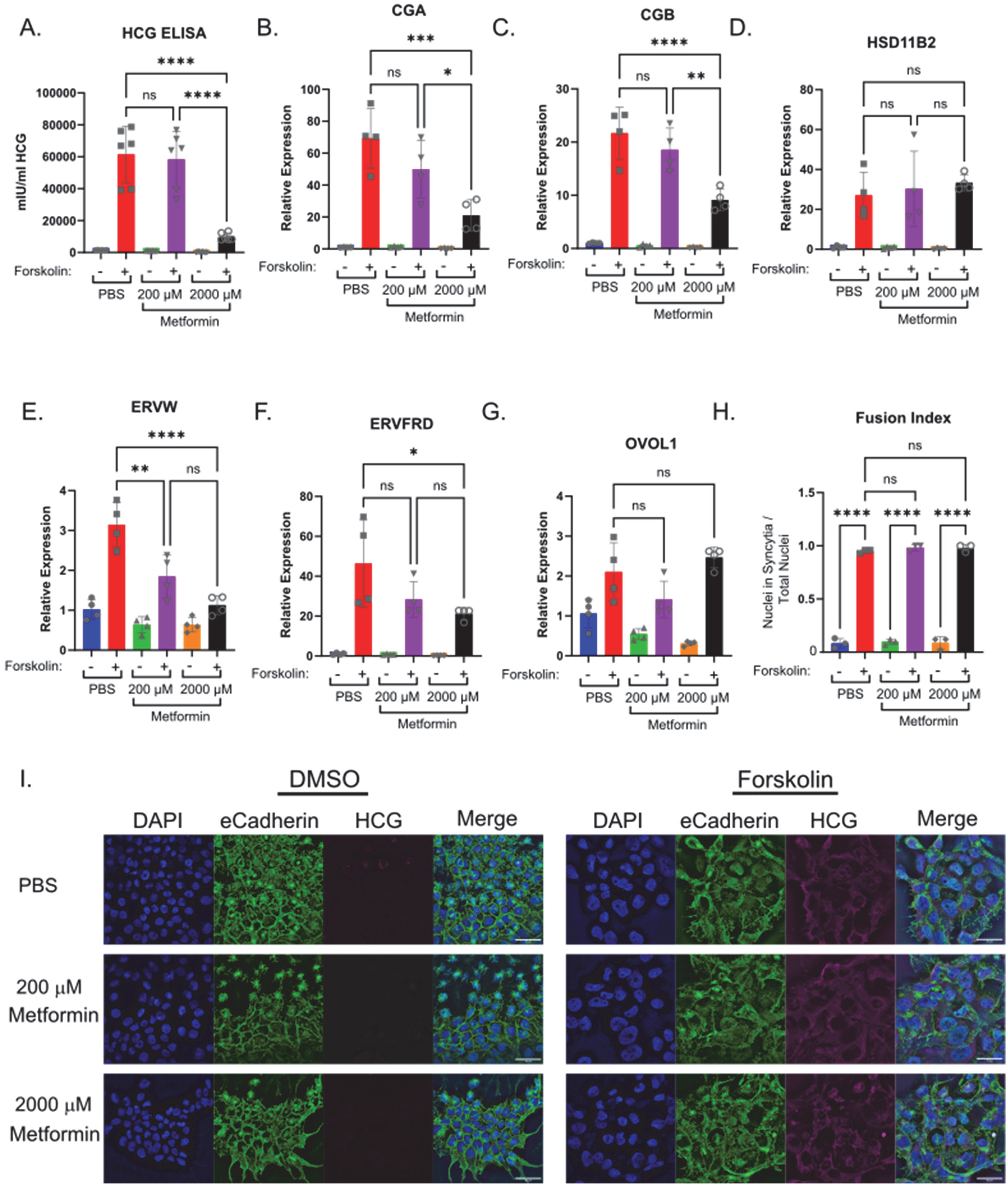
Increasing concentrations of metformin impair biochemical syncytialization. Forty-eight hours following treatment of BeWo cells with DMSO (0.4%) or 40 μM forskolin (FSK) in the presence of vehicle, Metformin 200 μM, or Metformin 2000 μM, we assessed: A) Production of HCG quantified by ELISA. n=6 biologic replicates. B-G) Relative gene expression by qPCR: B) CGA, C) CGB D) HSD11B2 E) EFVW F) ERVFRD G) OVOL1. n= 4 biological replicates. Data are representative of mean +/- SEM. I) Representative images of nuclear (DAPI, blue), cell membrane (E-cadherin, green), and HCG (magenta) stained BeWo cells treated for 48 hours with DMSO or forskolin (FSK) in the presence of vehicle, Metformin 200 μM, or Metformin 2000 μM. Scale bar represents 50 micron. H) Quantification of fusion index (number of nuclei in syncytia divided by total number of nuclei). n=6 biologic replicates and 3 fields of view from each replicate. Data are representative of the mean +/- SEM. *, *p*<0.05; **, *p*<0.01; ***, *p*<0.001; and ****, *p*<0.0001.

Trophoblast differentiation requires upregulation of genes necessary to support transcription, synthetic and fusogenic machinery during trophoblast differentiation (Costa, 2016;Renaud and Jeyarajah, 2022). To test the impact of metformin on this gene reprogramming, we treated BeWo cells with metformin and induced differentiation. In PBS treated cells, forskolin increased expression of CGA and CGB (involved in HCG production), HSD11B2 (regulator of glucocorticoid metabolism), ERVW and ERVFRD (fusogenic proteins), and OVOL1 (transcription factor) as expected **(Figure 4B-G).** Differentiation in the presence of 200 μM metformin similarly increased expression of CGA, CGB, HSD11B2, ERVFRD and OVOL1 **(Figure 4 C, D, F, G)**. However, ERVW expression upon forskolin differentiation was lower **(Figure 4E)** in presence of 200 μM of metformin in comparison to PBS. Treatment with 2000 μM metformin impaired forskolin induced upregulation of CGA, CGB, ERVW and ERVFRD in comparison to PBS **(Figure 4 B, C, E, F)**. However, no difference in OVOL1 or HSD11B2 expression was detected following 2000 μM metformin treatment compared to vehicle (**Figure 4 D, G**). This suggests that 2000 μM metformin may selectively impair transcriptional reprogramming of BeWo cells during differentiation.

To assess morphologic syncytialization, BeWo cells were plated on glass coverslips in the presence of vehicle, 200 or 2000 μM metformin prior to inducing syncytialization. 48 hours following differentiation, coverslips were stained with antibodies to HCG, which increases upon differentiation, and E-cadherin, which decreases upon differentiation as membrane fusion occurs (**Figure 4I**). The degree of fusion was quantified through the fusion index which identifies the number of nuclei found within syncytia divided by the total number of nuclei. We found robust induction of syncytialization upon treatment with forskolin with >90% of nuclei found in syncytiotrophoblasts in control cells. Treatment of BeWo cells with 200 and 2000 μM metformin resulted in similar degrees of syncytialization, with no statistically significant differences noted in fusion index **(Figure 4H).** Combined with gene expression data, this suggests that supra-therapeutic metformin treatment may de-couple biochemical and morphologic differentiation, which has been previously described (Cheng, 2004;Orendi et al., 2010;Leduc et al., 2012;Omata et al., 2013).

Given that differentiation regulates expression of nutrient transporters (Karahoda et al., 2022), we assessed changes in nutrient transporter expression in the presence of metformin. This may reflect a mechanism by which metformin could impact fetal growth. SNAT1 and SNAT2 **(Figure 5 A, B)** are involved in transport of neutral amino acids, and no difference in expression was detected following treatment with 200 or 2000 μM metformin. LAT1 and LAT2 **(Figure 5 C, D)** are involved in transport of L-amino acids, and following treatment with vehicle, 200 or 2000 μM metformin, a similar forskolin dependent increase in expression was observed in all investigated groups. Finally, we assessed expression of FATP6 **(Figure 5 E),** which is involved in transport of fatty acids, and observed increased expression following forskolin treatment in cells treated with 2000 μM metformin compared to vehicle control. This suggests that high doses of metformin may selectively impact nutrient transporter expression.

**Figure 5:**
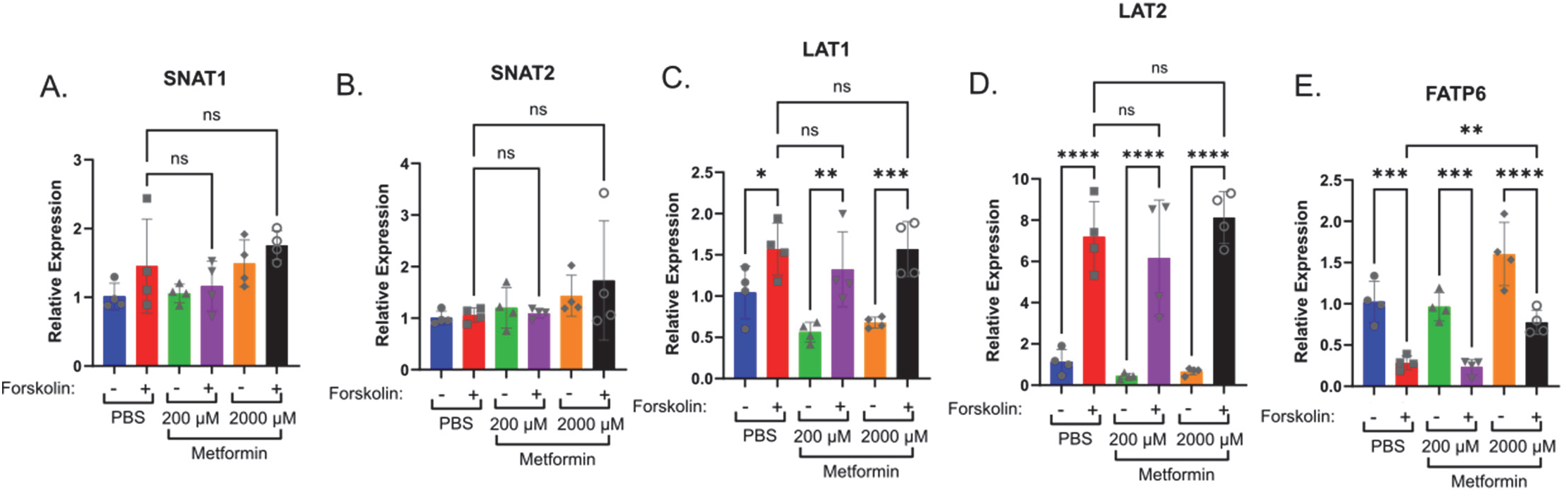
High doses of metformin may selectively impact nutrient transporter expression. A) SNAT1, B) SNAT2, C) LAT1, D) LAT2 and E) FATP6 gene expression by qPCR in BeWo cells treated with DMSO or Forskolin and 200μM or 2000μM metformin compared to vehicle. n= 4 biological replicates. Data are representative of mean +/- SEM. *, *p*<0.05; **, *p*<0.01; ***, *p*<0.001; and ****, *p*<0.0001.

### Impact of metformin on trophoblast stem cell model of differentiation

While Bewo cells are a widely accepted model of syncytialization, we sought to test the impact of metformin on trophoblast differentiation in another established model system. In the trophoblast stem cell model (Okae et al., 2018), cytotrophoblasts isolated from first trimester placenta are maintained in culture in the presence of WNT and EGF activators and TFG-B, histone deacetylase and ROCK inhibitors. Syncytialization is induced by removing these WNT, EGF, and HDAC modulators and adding a low dose of forskolin.

In the trophoblast stem cell model, 2000 μM metformin resulted in cell death 1in both self-renewing (SR) and differentiation (ST) conditions. However, we did not see evidence of cell death at the 200 μM metformin concentration. Comparing the impact of 200 μM metformin to vehicle, we did not detect any difference in HCG production by ELISA **(Figure 6 A).** No expression differences were observed in CGA, CGB, HSD11B2, ERVW, ERVFRD or TEAD4 following treatment with 200 μM metformin compared to vehicle (**Figure 6 B**). This again suggests that therapeutic concentrations of metformin do not dysregulate trophoblast differentiation.

**Figure 6:**
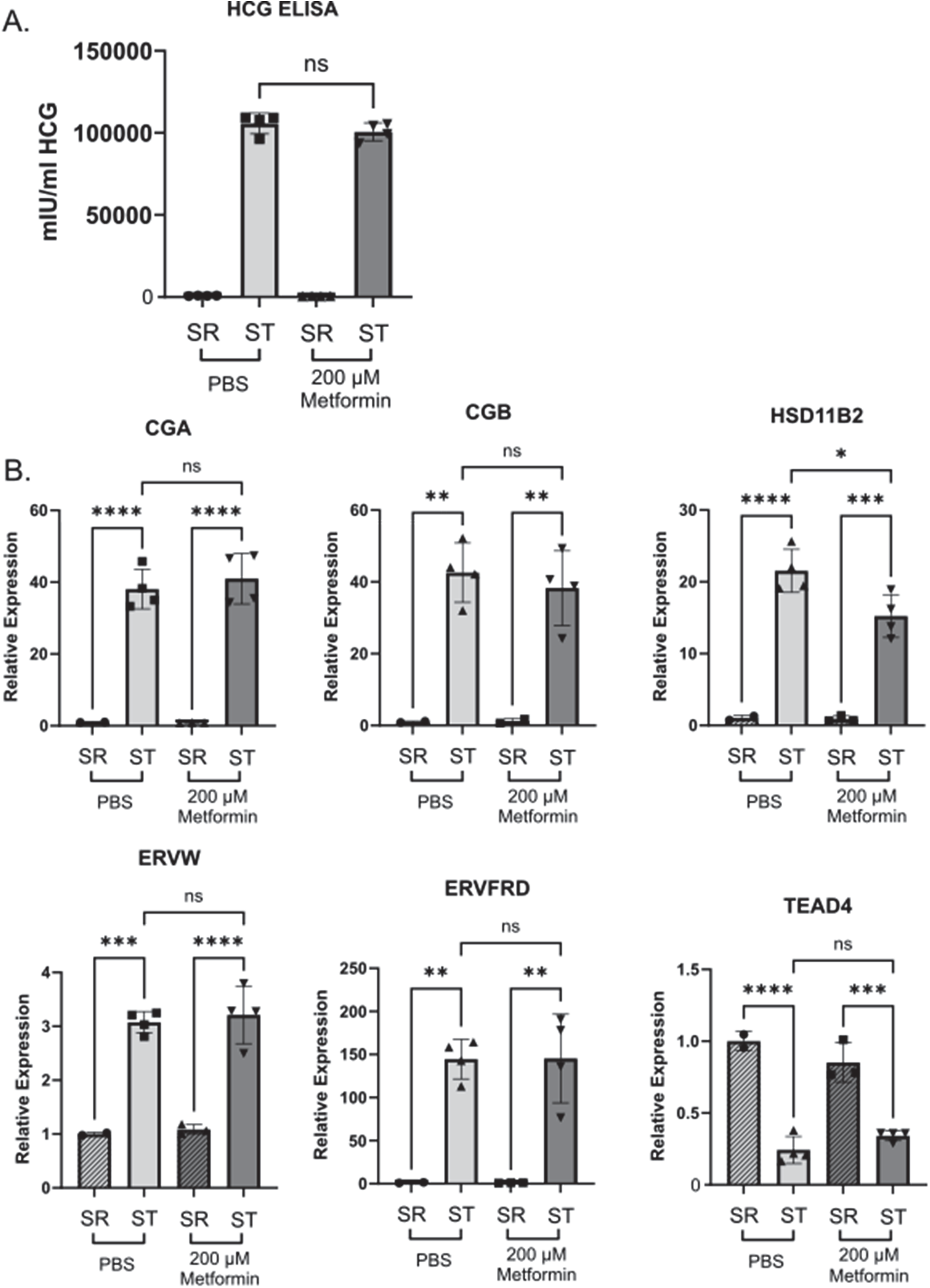
Metformin does not impair trophoblast stem cell differentiation. A) Production of HCG quantified by ELISA in self renewing (SR) cells or syncytiotrophoblast (ST) Trophoblast stem cells treated with vehicle (PBS) or 200 μM metformin. n=6 biologic replicates. B) CGA, CGB, HSD11B2, ERVW, ERVFRD, and TEAD4 gene expression by qPCR in SR or ST Trophoblast stem cells. n= 4 biological replicates. Data are representative of mean +/- SEM. *, *p*<0.05; **, *p*<0.01; ***, *p*<0.001; and ****, *p*<0.0001.

## Discussion

In this work, we evaluated the impact of metformin on trophoblast metabolism and differentiation using established cell culture models of trophoblast differentiation. We found that millimolar doses of metformin inhibited oxidative phosphorylation, resulting in increased glycolysis, lactate and TCA metabolites. Similar metabolic changes did not occur with 200 μM metformin treatment, consistent with prior reports that therapeutic concentrations of metformin may not function through impairment of Complex I activity. While treatment with 2000 μM of metformin impaired gene expression and HCG production following differentiation, 200 μM of metformin did not result in similar impairments. This overall suggests that therapeutic concentrations of metformin do not impair trophoblast differentiation.

Despite its widespread clinical use, metformin’s mechanism of action is not completely defined (LaMoia and Shulman, 2021;Aye et al., 2022). However, millimolar dosages are consistently associated with inhibition of Complex I, which results in activation of AMPK and its downstream pathways. Recent work examining the impact of 10 and 100 μM metformin on primary trophoblasts demonstrates statistically significant differences in basal respiration, maximal respiration and ATP production in primary trophoblasts treated with vehicle or 100 μM (Tarry-Adkins et al., 2022b). While this is a different cellular model system, the absolute magnitude of respiration differences remain small and are not normalized to account for variations in cell number. In our assay, we do not detect differences in cellular respiration at the 200 μM concentration, though profound impacts are seen at 2000 μM concentrations. Additionally, while we find broad changes in relative metabolite abundance with 2000 μM metformin treatments, we do not see similar changes between vehicle and 200 μM metformin treatments. Overall, this suggests that the potential benefits of metformin in pregnancy may not be attributable to large changes in trophoblast metabolism.

Prior studies have investigated metformin’s impact on osteogenic, neuronal, myogenic and adipogenic differentiation with varying results (Reviewed in (Jiang and Liu, 2020)). These reports highlight variability in metformin’s impact on differentiation across multiple system. This may be due to differences in metformin treatments or reflect inherent differences in differentiation across different cell populations. Given that GDM is associated with potential impairments in trophoblast differentiation, we examined metformin’s impact on trophoblast differentiation. While supra-therapeutic levels of metformin impair trophoblast biochemical differentiation in our BeWo cell model, we do not detect impairments in differentiation at therapeutic levels of metformin in either the BeWo or trophoblast stem cell models of differentiation. This suggests that metformin may not be directly contributing to differentiation defects seen in GDM.

The mechanisms underlying 2000 μM metformin’s impact on trophoblast differentiation are not known. Prior reports have implicated AMPK activation as a regulator of trophoblast differentiation with shRNA knock down of AMPK impairing trophoblast differentiation in the SM10 Trophoblast cell line models (Carey et al., 2014;Waker et al., 2017). It may be that higher doses of metformin modulate AMPK activity and impair differentiation in our system. Mitochondrial metabolism contributes to epigenetic regulation of gene expression (Dai et al., 2020), and it may be that impairments in mitochondrial function at high doses of metformin dysregulate epigenetic reprogramming during trophoblast differentiation. Human placental explants treated with 7 mM metformin demonstrated increased levels of Histone 3 Lysine 27 acetylation (Jiang et al., 2020). This highlights the potential role for supra-therapeutic concentrations of metformin to epigenetically regulate trophoblast differentiation.

We primarily relied on the BeWo model of cellular differentiation which may not fully recapitulate all aspects of trophoblast metabolism and differentiation. Given this limitation, we evaluated the effect of two different concentrations of metformin using trophoblast stem cells model. Although high dose metformin resulted in cell death, near-therapeutic concentrations metformin did not impair trophoblast differentiation, again supporting the observation that metformin is unlikely to impair trophoblast differentiation into syncytiotrophoblasts at therapeutic concentrations.

Overall, this work suggests that therapeutic concentrations of metformin are not likely to strongly impact trophoblast metabolism or differentiation. Additionally, it highlights the dose dependent impacts of metformin on cellular metabolism and differentiation events. The potential beneficial impacts of metformin seen clinically in GDM are likely occurring independent of inhibition of complex I and trophoblast differentiation into syncytiotrophoblasts.

## Acknowledgements

S.A.W. is supported by the Reproductive Scientist Development Program (RSDP) by the Eunice Kennedy Shriver National Institute of Child Health and Human Development (K12 HD000849) and the American Board of Obstetrics and Gynecology. M.D.G. is supported by 5R21HD102770. E.U.A. is supported by R56DK131447. P.A.C. is supported by R01DK091538 and R01AG069781. This work was supported by the resources and staff at the University of Minnesota Genomics Center (https://genomics.umn.edu), Informatics Institute, and Imaging Centers (SCR_020997).

## Author Contributions

S.K.N. and S.A.W. conceived of the study, designed and completed experiments, analyzed data, prepared figures, and wrote the manuscript. R.M.M. performed experiments, analyzed data, and prepared figures. S.J. and D.S. performed experiments. A.B.N., P.A.C., and P.P. assisted with mass spectrometry analyses and interpretation. S.J. and E.U.A. performed Seahorse experiments, analyzed, and interpreted data. M.D.G. assisted with TSC experiments and data interpretation. All authors reviewed the manuscript.

## Key Resources

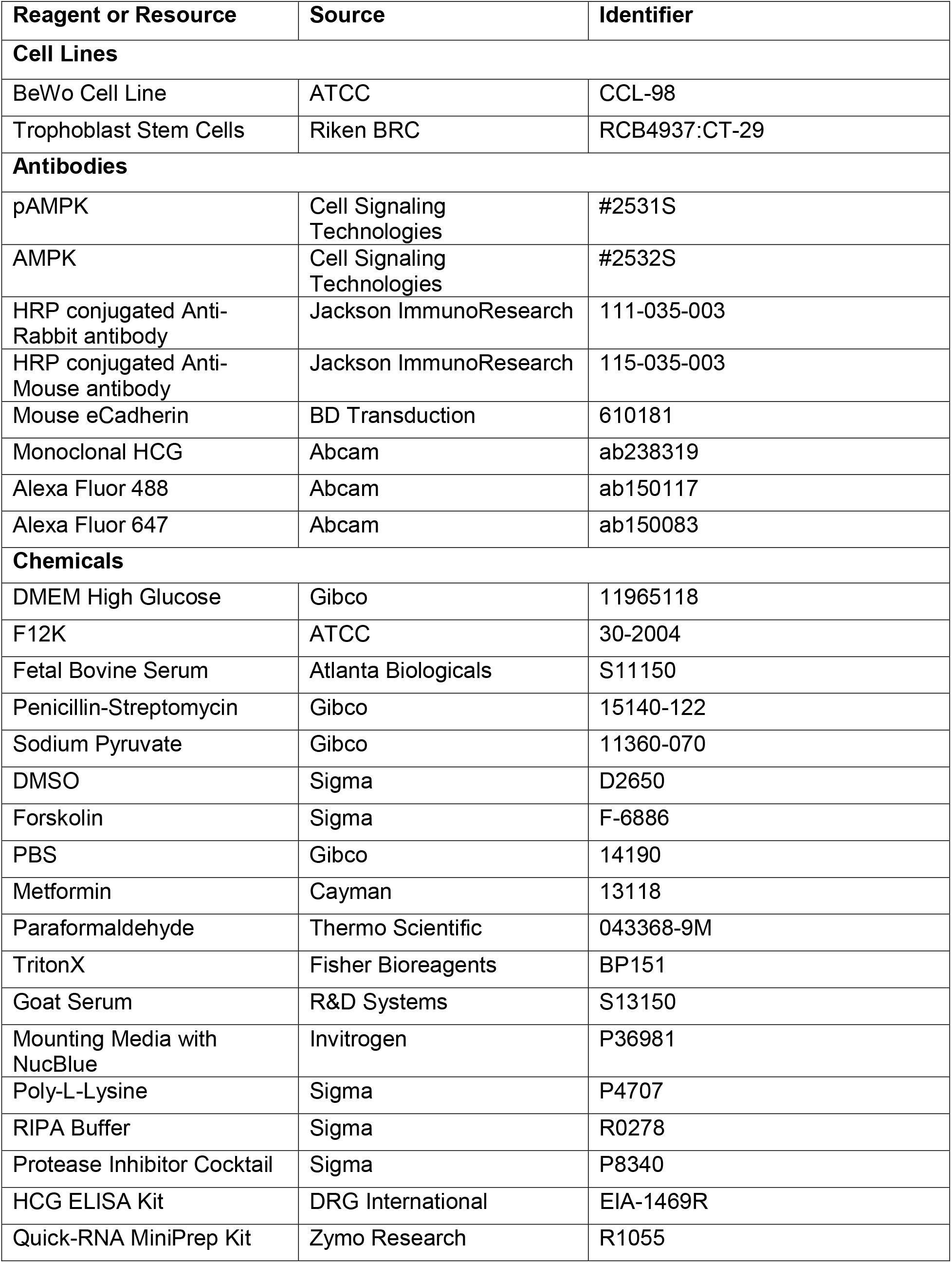
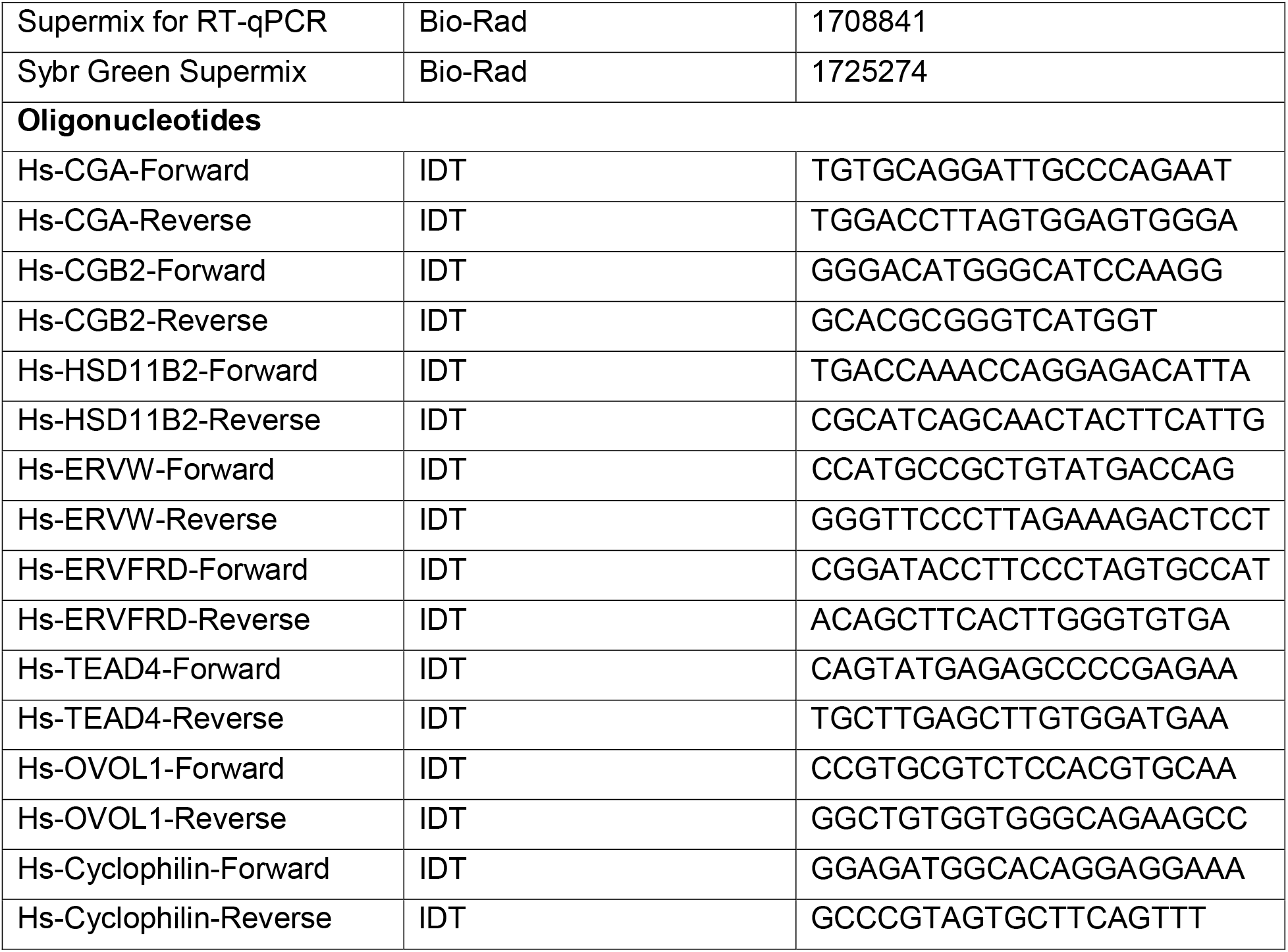

