## Supplemental Figure for "Metformin impairs trophoblast metabolism and differentiation in dose dependent manner"

**Supplemental Figure 1: Metformin impacts extracellular acidification rates**

A-C) Normalized extracellular acidification rate for BeWo cells treated with vehicle, 200 µM metformin, or 2000 µM metformin the presence of DMSO (0.4%, vehicle) or 40 µM forsksolin (FSK) for 48 hours. n= 4 independent replicates per condition

D) Basal ECAR for BeWo cells treated with vehicle, 200 µM metformin, or 2000 µM metformin. Data representative of mean +/- SEM. n=4 biologic replicates per condition. *, *p*<0.05; **, *p*<0.01; ***, *p*<0.001; and ****, *p*<0.0001.

**Supplemental Figure 1:**

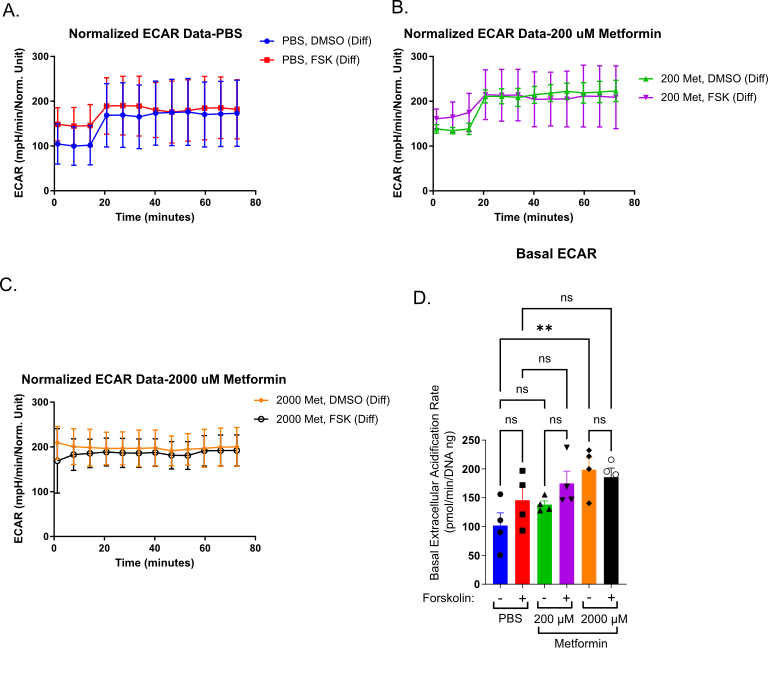
